# Predicting butyrate- and propionate-forming bacteria of gut microbiota from sequencing data

**DOI:** 10.1101/2022.03.06.483156

**Authors:** Berenike Kircher, Sabrina Woltemate, Frank Gutzki, Dirk Schlüter, Robert Geffers, Heike Bähre, Marius Vital

## Abstract

**Background:** The bacteria-derived short chain fatty acids (SCFAs) butyrate and propionate play important (distinct) roles in health and disease and understanding the ecology of respective bacteria on a community-wide level is a top priority in microbiome research. The aim of this study was to reveal members harboring main pathways for the production of those metabolites and assess the applicability of sequence data (metagenomics and 16S rRNA gene) to predict SCFAs production *in vitro* and *in vivo*.

**Results:** A clear split between butyrate- and propionate-forming bacteria was detected with only very few taxa exhibiting pathways for the production of both SCFAs. After *in vitro* growth of fecal communities from distinct donors (*n*=8) on different substrates (*n*=7) abundances of bacteria exhibiting pathways correlated with respective SCFA concentrations, in particular in the case of butyrate. While final growth differed markedly between cultures, communities showed high functional redundancies with comparable yields, i.e., concentration of metabolite per grown bacterium exhibiting pathway(s), irrespective of the donor and substrate used. For propionate, correlations were weaker indicating that its production is less imprinted into the core metabolism compared with butyrate-forming bacteria. Longitudinal measurements *in vivo* (five samples derived from 20 subjects) also revealed a correlation between abundances of pathway-carrying bacteria and concentrations of the two SCFAs. Additionally, lower bacterial cell concentrations, together with higher stool moisture, promoted overall bacterial activity (measured by flow cytometry and coverage patterns of metagenome-assembled genomes) that led to elevated SCFAs concentrations with over-proportional levels of butyrate. Butyrate concentrations displayed lower temporal stability than propionate, however, abundances of bacteria exhibiting the butyrate-forming pathway were more stable than those carrying pathways for propionate production. Predictions on pathway abundances based on 16S rRNA gene data using our in-house database worked well yielding similar results as metagenomic-based analyses.

**Conclusions:** We demonstrated that pathway abundances enable predictions on concentrations of SCFAs indicating that stimulating bacterial growth directly leads to more production of those compounds. The strong separation of gut microbiota into two functional communities facilitates the development of precision intervention strategies targeting either metabolite.

## Introduction

Short-chain fatty acids (SCFAs), mainly acetate, butyrate and propionate, are major products of bacterial fermentation in the human large intestine and have increasingly become the focus of research due to their importance for host metabolism and health. They are known to reduce local and systemic inflammation processes by immunomodulatory properties and maintenance of gut epithelial integrity (1–3). Scarcities of SCFAs are associated with emerging noncommunicable metabolic disorders, such as cardiovascular disease, obesity and type II diabetes (4), and an impairment of colonization resistance against enteric pathogens (5)(6). Despite common actions of SCFAs, they also markedly differ in their effects on the human body. For instance, the main target site of butyrate are colonocytes that use this compound for energy generation, whereas the bulk of propionate reaches the liver and promotes gluconeogenesis (7). Circulating SCFAs bind to G-protein-coupled receptors that are expressed throughout the body, however, SCFA affinities to individual receptor types differ (1) promoting distinct levels of response (8). Furthermore, only butyrate shows epigenetic properties that plays a role in diverse diseases (9)(10).

Direct measurements of fecal SCFAs represent the gold standard for assessing a given community’s capability to produce those compounds. However, SCFAs are volatile molecules demanding immediate preparation of samples for exact measurements, which is often difficult in practice. Furthermore, it is estimated that 90 - 95 % of SCFAs produced are absorbed by the colonic epithelial and inferences on production of individual SCFAs from measured concentrations is, hence, doubtful (11, 12).

While acetate is produced by most members of gut microbiota, only specific, phylogenetically distinct bacterial groups form butyrate and propionate (13). Butyrate formation from carbohydrates is performed via the Acetyl-CoA pathway (14), whereas propionate is largely produced via the succinate and propanediol pathways that act on carbohydrates as well (15). The former two pathways are anchored in the core metabolism of bacteria making them essential biochemical routes for growth of respective bacteria. It is expected that the propanediol pathway is also important for bacteria to occupy niches *in vivo* (15), whereas several protein-fed pathways leading to the formation of butyrate are not essential for bacteria to grow in the gut and are believed to play minor roles (16). A systematic screening of (meta)genomes for exhibiting butyrate synthesis pathways in gut microbiota has been performed demonstrating that primarily members of the *Lachnospiraceae* and *Ruminococcaceae* of the *Firmicutes* serve as butyrate producers. In case of propionate, the succinate pathway is suspected to be predominantly encoded on gut bacteria of the phylum *Bacteroidetes*, including the abundant *Bacteroides*, and a few *Negativicutes* of the *Firmicutes*, whereas propanediol pathway carriers almost exclusively belong to the *Lachnospiraceae*, mainly from the genera *Ruminoccus* and *Blautia* (15). However, a systematic screening for those pathways in genomes derived from the gut environment is lacking.

A major goal in gut microbiota research is to get (quantitative) insights into bacterial functions affecting host physiology. In this context, deciphering contributions of individual bacteria of a given community to the total SCFA pool is a top priority (17). While metagenomic data allow for exact determination of SCFA pathway distributions in a given sample (16), analyses are often tedious and inferring functionality from low-cost, high-throughput data such as 16S rRNA gene results is desirable. An additional aspect that has increasingly become recognized is bacterial load, as it was demonstrated that cell numbers per gram stool differ by an order of magnitude, which probably has profound influences on overall functionality in quantitative terms as well as actual metabolite concentrations of a given sample (18).

The aim of this study was to reveal butyrate-and propionate forming communities of gut microbiota in quantitative terms and assess the ability to predict the production of those two SCFAs based on sequence data. To this end, we comprehensively screened reference organisms of gut bacteria for exhibiting respective pathways and performed a series of *in vitro* incubations together with a longitudinal *in vivo* experiment including human subjects, where a multitude of parameters considered important for SCFAs production were analysed.

## Materials and methods

### *In vitro* incubation experiments

Stool from eight healthy subjects (5 female/3 male), that were also participating in the *in vivo* study below, was collected and immediately transferred to a vinyl anaerobic chamber (Coy Laboratory Products, Grass Lake, MI, USA) for experiments. Samples were diluted (1:100) in preproduced 1x PBS, subjected to 30 µm-filtration (Miltenyi Biotec, Bergisch Gladbach, Germany) and added to anaerobic basal medium (described in (19), excluding bile acids) to achieve a starting concentration of ∼1-3 × 10^7^ cells mL^−1^ (an aliquot for enumerating cell concentrations by flow cytometry (see below) was diluted fivefold in 1x PBS, snap frozen in liquid nitrogen and stored at -20 °C). Suspensions were aliquoted (10 ml) into Hungate tubes and 1 ml of individual growth substrates were added (final concentration of 2 g L^−1^). The following growth substrates were used: resistant starch type 2 and 3 (Hylon VII (PCR) and Novelose330; both from (Ingredion, Manchester, UK)), pectin from apple (Sigma Aldrich, St.Louis, MO, USA), mucin (Sigma Aldrich, St.Louis, MO, USA), inulin (Orafti HP; from Beneo-Orafti, Oreye, Belgium) and protein (Bacto Casitone, BD, Franklin Lakes, NJ); a negative control (stool in basal medium) was included as well. Substrates were boiled for 5 min in a microwave and pre-reduced overnight under the anaerobic chamber before being used in the experiments. Incubations were carried-out in duplicate samples at 37°C for 24 h (200 rpm). The pH was determined using a pH-Meter (Knick International, Berlin, Germany) with and Inlab-semi-micro electrode (Mettler Toledo, Columbus, Ohio, USA). 2 mL of cultures were centrifuged (15.400 g, 4 °C), diluted in NaOH (5 mM) and stored at -80 °C before determination of SCFA concentrations (see below); the pellet was used for DNA extraction. Bacterial concentrations were determined by flow cytometry (see below).

### Monitoring of gut microbiota *in vivo*

Twenty volunteers (11 female/9 male) provided 5 fresh stool samples over a period of 3 months; 3 samples were collected in November/Dezember 2019 (2 weeks interval), whereas another 2 samples (2 weeks interval) were collected in January 2020. Approximately 2 g stool were collected into faeces collection tubes (Sarstedt, Nümbrecht, Germany), put at 4 °C and diluted fivefold in 1 x PBS; undiluted aliquots (∼200 mg) for DNA extraction were snap frozen in liquid nitrogen and stored at -80°C until further analysis. For flow cytometric analyses an aliquot of 100 µl from the dilution was snap frozen, whereas for the determination of SCFAs concentrations 20 µL were added to 980 µL NaOH (5 mM), centrifuged (5 min, 4500 g, 4°C) and 100 µL of the supernatant was collected in gaschromatography (GC) glass vials (Macherey-Nagel, Düren, Germany); both were stored at -80°C. Faecal pH was directly measured in stool suspensions (fivefold dilution in distilled water). The bristol stool scale (BSS) was recorded by individual donors themselves and determinations of dry weight was performed by weighing aliquots of approximately 200 mg stool before and after drying via SpeedVac RVC 2-18 CD plus (Martin Christ Gefriertocknungsanlagen, Osterode am Harz, Germany), at 37 °C (1300 rpm for 4 h). The study was approved by local ethic authorities (#8566_BO_K_2019) and all subjects have given informed consent.

### Flow-cytometric analyses and determination of SCFA concentrations

For flow-cytometric measurements (FCM) the fivefold dilutions of stool samples (*in vivo* experiments) were thawed at room temperature and diluted 100x with 1x PBS, including a 30 µm-filtration step (Miltenyi Biotec, Bergisch Gladbach, Germnay). For *in vitro* samples, 1:500 dilutions (1x PBS) were directly prepared from growth cultures; samples taken at the beginning of the experiment were thawed. All suspensions were stained with 10 µl EDTA and 10 µl SYBR Green (Thermo Fisher Scientific, United States) according to (20) and incubated for 15 min at 37 °C in the dark. Before measurements, stained samples were diluted 10-fold in 1x PBS and cell concentrations as well as green fluorescence intensities were recorded on a MACSQuant Analyzer 10 (Miltenyi Biotec, Germany).

Concentrations of acetate, butyrate and propionate of faecal samples (*in vivo* experiments) and *in vitro* incubations were quantified at the RCU Metabolomics of Hannover Medical School based on a GC-MS method including a derivatization step and addition of a labelled standard (see Supplemental Methods).

### DNA extraction, library preparation and sequencing

DNA was extracted (DNeasy PowerSoil Pro Kit, Qiagen, Germany; including a beat-beating step (2×20sec on Fastprep System (MP Biomedicals, Santa Ana; California; USA) at speed 5.5)) and libraries for shotgun-sequencing were prepared (Illumina DNA Prep, Illumina, United States) that were subsequently sequenced on Illumina NovaSeq 6000 (at Helmholtz Centre for Infection Research (HZI)) in paired-end mode (2 × 150 bp). For *in vivo* experiments 2 × 10^7^ reads per sample were sequenced (for the first and last samples 5 × 10^7^ (2 × 250 bp) were obtained), whereas shallower sequencing (5 × 10^6^ reads) were performed for *in vitro* samples. Libraries for 16S rRNA gene sequencing were prepared according to (21), but targeting the V3V4 region using primers from (22) with an annealing temperature of 55 °C. Obtained amplicons were pooled and sequenced on Illumina MiSeq (2 × 300 bp).

### Constructing the catalogue of SCFA pathway genes

All representative genomes from the Unified Human Gastrointestinal Genome (UHGG) collection (23), that displayed decent quality (completeness >80 and contamination <10) were included into analysis (n=3,207). To increase diversity, high quality isolates (completeness >95%, contamination <2% and a 16SrRNA gene length >70 %) that showed an average nucleotide identity (ANI, determined via fastANI (v1.32) (24)) below 98% to the representative (and to each other) were included as well (n=522); A few were manually selected (n=25). Genomes were downloaded and gene sequences were extracted with GffRead (25). UBCG (v3.0) was used to make a phylogenetic tree based on 92 house-keeping genes (HKGs) (26) and genomes were screened for SCFA pathways. For butyrate, the same approach as described previously was used (16) consisting of a multi-level approach involving Hidden Markov Models (HMM) for all genes of the Acetyl-CoA (ACoA) pathway and analyses on gene-synteny and pathway completeness. For propionate, a similar multilevel-screening approach for the succinate (Suc) and propanediol (Pdiol) pathways based on key genes defined by Reichardt et al (2014) (15) was developed. Details are given in the Supplemental Methods. All results of pathway screenings were manually checked and a few genomes, that were filtered-out due to fragmented pathway genes, were included. For annotations based on the Kyoto Encyclopedia of Genes and Genomes (KEGG), all genomes were subjected to GhostKOALA (27) and subsequent filtering based on key genes of individual pathways (for ACoA only *bhbd, cro* and *bcd* were considered) was performed including manual inspections (e.g. lacking of single genes). For 16S rRNA gene analyses, 1,623 genomes that had decent length genes (>900bp) were included. The longest gene from each genome were aligned in Clustal Omega (webserver) and a phylogenetic tree was constructed via FastTree 2 (v2.1.10) (28). Manual inspections led to removal of 24 sequences as they clustered with the wrong phylum. Duplicates were removed and finally 1,556 16S rRNA gene sequences were used for follow-up analyses.

### Metagenomic analyses

The metaWRAP pipeline (v1.3.0) was used for genome-resolved metagenomic analyses (29). Raw sequences of *in vivo* samples were quality filtered, assembled via MEGAHIT and binned via a combination of MaxBin 2, MetaBAT 2 and CONCOCT using the BIN_REFINEMENT module; for assembly, all samples of a person were merged, whereas they were treated separately during the binning process. Finally, bins were reassembled (REASSEMBLE_BINs module) yielding metagenome-assemble genomes (MAGs). Taxonomic annotations of MAGs were done via the GTDB-Tk (1.7.0) (30), HKGs were extracted via UBCG and SCFA-forming pathways were detected as described above. GRiD (v1.3.0) was used to infer growth rates by calculating coverage ratios between *ori* and *ter* (31); a cumulative value for each sample was calculated (average of all MAGs normalized for their relative abundances).

For determining SCFAs-pathway abundances, HKGs and pathway genes from UHGG references and from constructed MAGs were used as a catalogue (non-target pathway genes showing sequence similarity, but displayed HMM scores below the set cut-offs, were included as well, as described previously (16)) for mapping reads via BBmap2. Resulting counts were gene-length corrected and normalized to HKGs (mean abundance of all HKGs) yielding relative abundances of genes from respective pathways; mean results of pathway genes were used in follow-up analyses as done previously (16). Taxonomic affiliations of individual pathways were determined on the species level, where a taxon was recorded if at least 3 genes of a pathway were detected (2 genes for the Pdiol pathway), which included 95.5 % +/- 2.7 % (*in vivo*) and 91.3 % +/- 8.3 % (*in vitro*) of all reads mapped to pathway genes. Results of taxonomies were subsequently merged for insights at higher orders. For overall taxonomic compositions HKGs were used, where a species was considered present if at least 20 HKGs were detected (96.4 % +/- 1.0 % (*in vivo*) and 92.5 % +/- 2.5 % (*in vitro*) of all reads that mapped to HKGs were included).

### 16S rRNA gene analyses

Sequences were processed via the DADA2 pipeline (v1.20) in default mode and annotated based on RDP’s taxonomy (32). Chimera were removed and only sequences displaying a length >300bp, counts >10 and annotated at the phylum level were included (sequences derived from Chloroplasts were excluded). SCFAs pathway predictions were done via the picrust2 algorithm (v2.3.0b) (33) by placing sequences into our reference tree (*place_seqs*.*py*) followed by hidden-state predictions (*hsp*.*py*).

### Statistics and generation of plots

Growth of SCFA-pathway exhibiting bacteria was calculated from final cell concentrations determined by FCM and metagenomic results that provided relative abundances of bacteria carrying individual pathways. All plots were constructed in R via ggplot2 (v3.3.5) and *ggtree* (v1.14.6). Correlations for *in vitro* results were calculated (function *lm*) from original and log-transformed data (log(data+1)). Non-metric multidimensional scaling (NMDS) analyses were done in *phyloseq* (v1.36.0) on relative abundance data of taxa on the species level. Correlations of *in vivo* parameters were determined via linear mixed-effects models using the function *lmer* from the lme4 package (v1.1-27.1) on log-transformed data (log(data+1)) including subject as a random effect.

## Results

### Establishing a database of gut bacteria harbouring major pathways for the production of butyrate and propionate

We screened in total 3,754 genomes, involving 3,207 species representative genomes, originating from the UHGG collection for exhibiting butyrate and propionate-forming pathways. In total, 18.0 % (n=675) of genomes were classified as butyrate producers harbouring the ACoA pathway, while 14.9 % (*n*=558) and 9.3 % (*n*=350) were exhibiting the Suc and Pdiol pathways for propionate synthesis, respectively (Figure 1). For the former pathway, 50.9 % carried butyryl-CoA acetate CoA-transferase (*but*) as the terminal enzyme. Butyrate kinase (*buk*) was detected on 20.9 % of genomes and 1.2 % (all members of the genus *Coprococcus*) exhibited both enzymes, whereas in 27 % of cases neither gene was detected. While pathways were present on a wide range of distinct taxa, the distribution of both the ACoA and the Suc pathway was largely consistent on the genus level. For instance, almost all members of the key butyrate-producing genera *Faecalibacterium* and *Agathobacter* exhibited the ACoA pathway and most members of the *Bacteroides* and *Phocaeicola* displayed the Suc pathway (Figure S1). Only a few metagenome-assembled genomes (MAGs) within those genera were predicted lacking respective pathways. The Pdiol pathway on the other hand clustered less homogeneously, however, members of several abundant genera of gut microbiota, such as *Blautia_A*, consistently exhibited this pathway. Overall, results suggest that main pathways are largely split between bacterial groups, where genomes either contained genes for the formation of butyrate or propionate. Of the 675 genomes harbouring the ACoA pathway only 8.6 % and 10.8 % also exhibited the Suc and Pdiol pathway, respectively. This functional division into butyrate- and propionate-forming communities was even more pronounced in *in vitro* and *in vivo* communities (see below).

**Figure 1.**
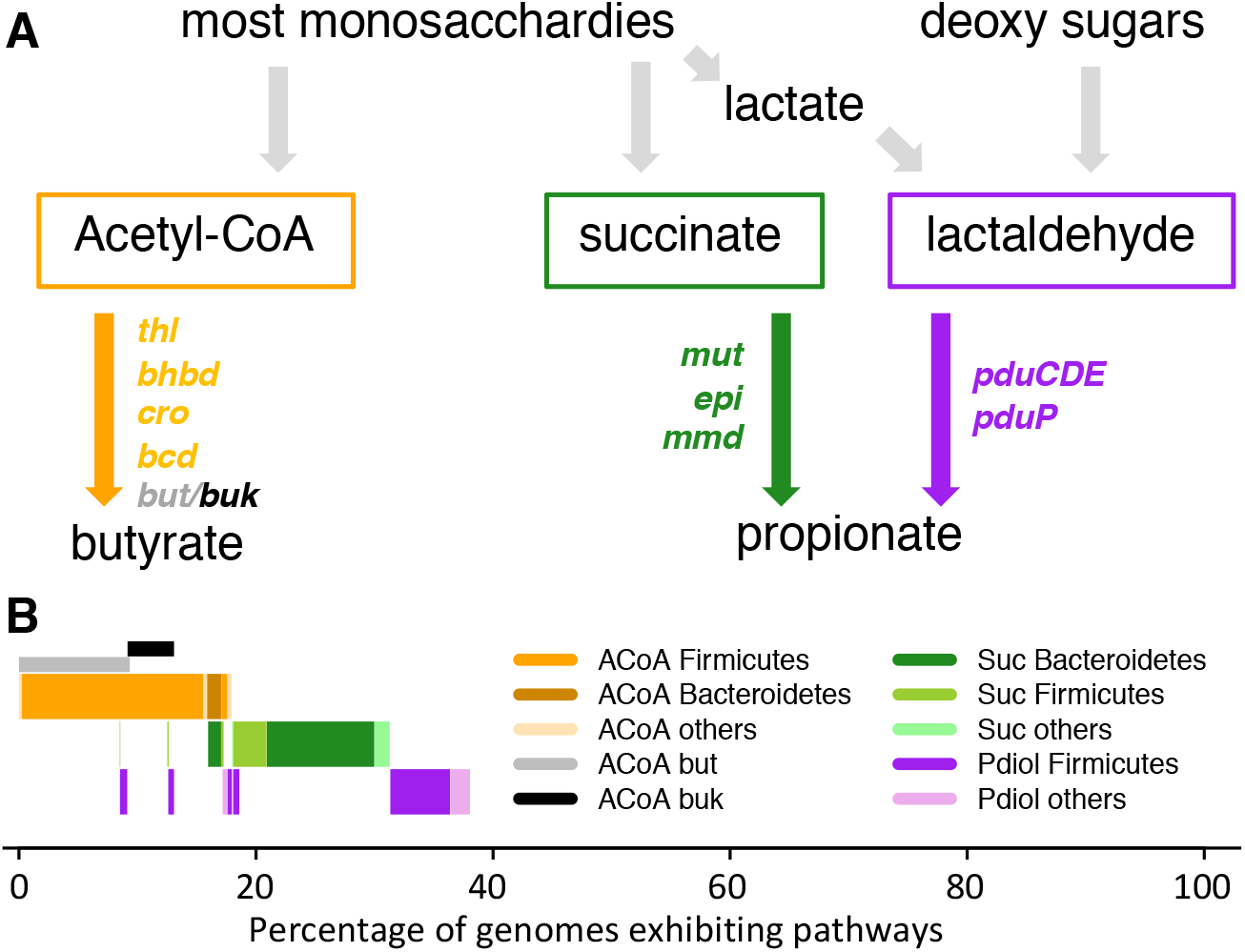
Overview of pathways and results from our constructed database. Panel A shows a simplification of main pathways involved in the formation of butyrate and propionate including gene names encoding enzymes catalysing individual steps. For detailed description of pathways see reference (13). Genomes of individual species of the Unified Human Gastrointestinal Genome (UHGG) collection were screened for exhibiting those pathways and taxonomic affiliations on the phylum level are indicated (panel B). ACoA: main butyrate-forming pathway including acetyl-CoA acetyltransferase (*thl*) β-hydroxybutyryl-CoA dehydrogenase (*bhbd*); crotonase (*cro*); butyryl-CoA dehydrogenase (*bcd*) as well as genes encoding the terminal enzymes butyryl-CoA:acetate CoA transferase (*but*) and butyrate kinase (*buk*). Suc: main propionate-forming pathway from carbohydrates including methylmalonyl-CoA mutase (*mut*), methylmalonyl-CoA epimerase (*epi*) and methylmalonyl-CoA decarboxylase (*mmd*). Pdiol: additional propionate-forming pathway with the key enzymes propanediol dehydratase (*pduCDE*) and propionaldehyde dehydrogenase (*pduP*).

Inferring pathways from genomes based on annotations derived from KEGG showed several discrepancies compared with our in-house database. Most obvious was the prediction of the ACoA pathway on genomes of many members of the *Proteobacteria*, such as *Acinetobacter spp*., *Aeromonas spp*., *Citrobacter spp*. and *Yersinia spp*. and several *Bacilli*, that have not been described as butyrate producers (Figure S1). Also for propionate-forming bacteria inconsistencies with KEGG were detected. For instance, KEGG suggested that a specific clade of the *Verrucomicrobiota* including *Akkermansia spp*., that are known propionate producer, lacks the Suc pathway. Furthermore, based on KEEG only a few members of the genus *Blautia_A* exhibit the Pdiol pathway, whereas our data indicate that it was highly prevalent in this genus (Figure S1).

Of the total 3,754 genomes analysed 41.4 % (n=1,556) exhibited high-quality 16S rRNA gene sequences and were used as references for predicting SCFAs pathways based on the *picrust* algorithm (Figure S2). In particular, many MAGs were devoid of adequate sequences and could, hence, not be included. Overall, predictions were largely following reference data, especially for the ACoA and Suc pathways and their presence/absence was wrongly predicted for only a few genomes. The Pdiol pathway was predicted correctly for most genomes as well, however, for a few taxa that disparately exhibit this pathway, such as *Enterocloster* and *Escherichia*, predictions deviated from references. Predictions based on input sequences trimmed to the variable regions V3V4 were largely mirroring full-length gene results (Figure S2).

### Incubations of gut communities *in vitro*

To investigate the predictability of sequence data for the production of butyrate and propionate, we conducted a series of *in vitro* experiments, where freshly provided stool samples, derived from eight individuals, were incubated with six different growth substrates, namely, resistant starches type 2 and type 3, pectin from apple, mucin, inulin, and protein. After 24 h, bacterial growth, i.e., cell numbers grown (measured by flow-cytometric (FCM) analysis), relative pathway abundances based on metagenomic analyses, and SCFAs concentrations (acetate, butyrate and propionate) were determined. Overall, bacterial composition after *in vitro* growth was comprised of common gut bacteria (Figure 2) and was similar to *in vivo* communities (see below). On average, we detected growth of 4.07×10^8^ ± 2.72×10^8^ mL^−1^ butyrate producers, i.e. bacteria exhibiting the ACoA pathway, comprising 32.6 % ± 6.9 % of the total community, while 2.95×10^8^ ± 1.62×10^8^ (mL^−1^) Suc and 1.44×10^8^ ± 1.00×10^8^ (mL^−1^) Pdiol pathway carrying bacteria were detected representing 27.8 % ± 12.8 % and 10.9 % ± 5.0 % of the overall community, respectively. A clear split between butyrate- and propionate-producers was observed and only 2.77 % of bacteria (mainly *Anaerobutyricum*) harboured pathways for both butyrate and propionate synthesis (Figure 2). The butyrate-forming community was primarily composed of *Firmicutes*, with several main genera including *Faecalibacterium* (27.7 %) and *Agathobacter* (10.2 %), whereas members of the *Bacteroidetes*, primarily *Bacteroides* (37.7 %) and *Phocaeicola* (33.6 %) comprised the Suc pathway; *Blautia_A* (44.6 %) and *Anaerobuytricum* (18.3 %) of the *Firmicutes* were the main members carrying the Pdiol pathway (Figure 2).

**Figure 2.**
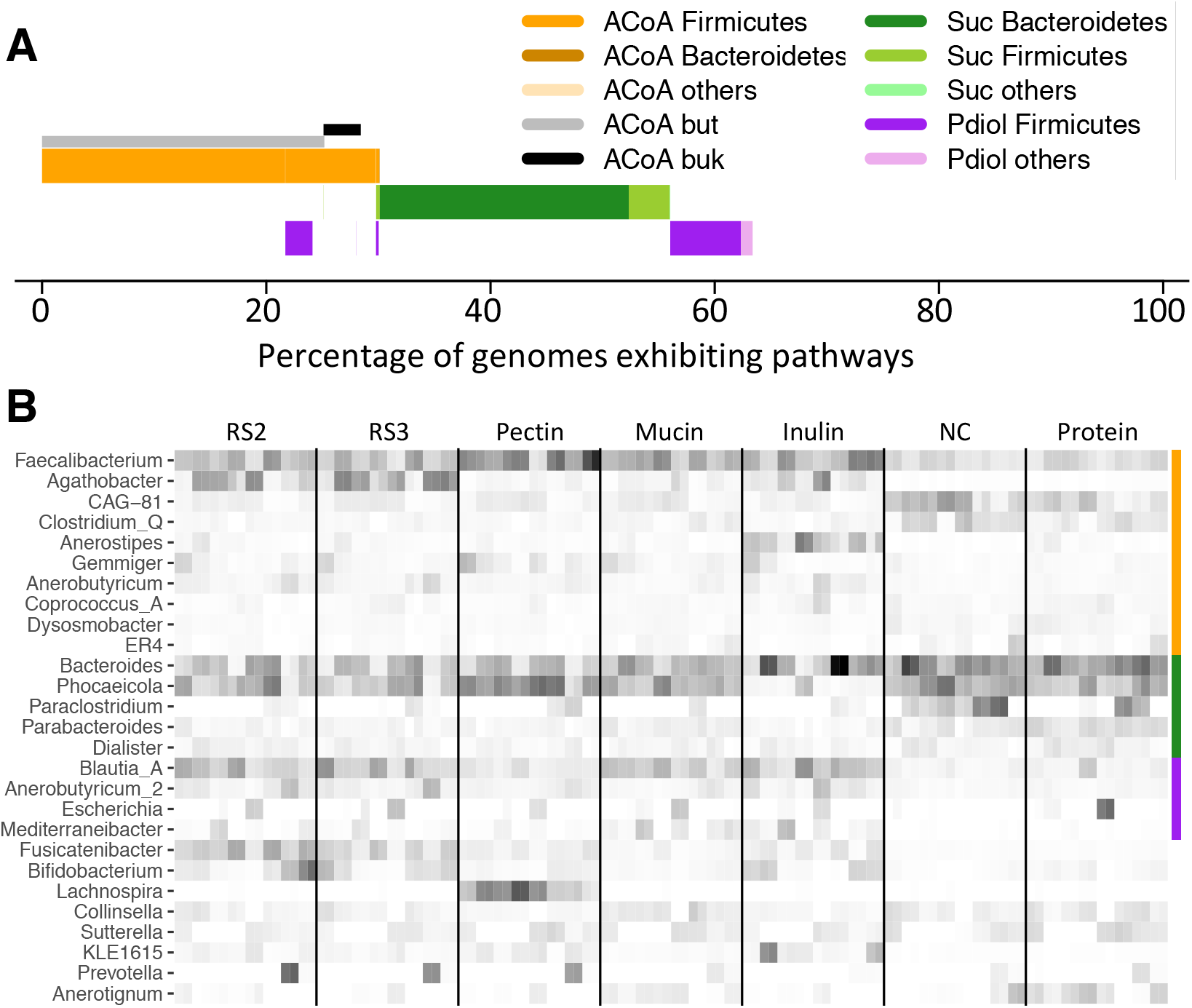
Overview of pathway abundances and associated taxonomic composition after *in vitro* growth of faecal communities derived from eight subjects grown on different substrates (n=7)). Panel A gives average abundances of pathways including taxonomic affiliations on the phylum level, whereas abundances of major genera comprising individual pathway communities (indicated by the colour bar on the right) are given in Panel B. Abundances are relative to house-keeping genes of the total bacterial community. Incubations were performed in duplicate samples. For abbreviations of pathways in panel A see Figure 1. RS2/3 (resistant starch type 2/3); NC (basal medium).

Global community structures on the species level clustered strongly according to their donors and communities growing on proteins and only the baseline medium showed unique patterns forming a separate group (Figure S3A). Composition of functional communities, i.e., taxonomic affiliation of individual pathway carriers, showed high subject-specific signatures as well (Figure S3B, C).

The average concentration of butyrate formed in all incubations was 5.27 mM ± 2.63 mM (Figure 3A) and we found a strong correlation (R^2^ = 0.63) with final growth of bacteria exhibiting the ACoA pathway. Relative butyrate concentrations (percentage of total SCFAs) was related with abundances of respective bacteria as well (R^2^ = 0.30; Figure 3B). Overall, (relative) butyrate production and yields, i.e. butyrate produced per cell harbouring the ACoA pathway, were in a similar range for all communities derived from the different subjects; samples inoculated with bacteria from two subjects (e and h) showed lower relative production and their yields were increased (Figure 3C-E). Growth on inulin and the resistant starches resulted in higher butyrate concentrations compared with results from mucin and pectin, whereas values for growth on proteins and the basal medium were the lowest (Figure 3F-H). For propionate, a positive correlation between grown bacteria that contain the Suc and Pdiol pathways and propionate concentrations was observed (R^2^=0.24; Figure 4A). The fraction of propionate of total SCFAs was, however, not associated with abundances of those bacteria (Figure 4B). (Relative) production was similar between communities derived from different subjects (Figure 4C-E) and average concentrations of 3.97 mM ± 1.35 mM were lower than those of butyrate. Most propionate was formed during growth with mucin compared with other substrates, whereas the yield was highest on the basal medium (Figure 4F-H). The yield for propionate was lower, namely 10.9 ± 5.4 fmol propionate per propionate producer, compared with that of butyrate-producing bacteria (15.5 ± 6.4 fmol butyrate per butyrate producer). Total SCFAs concentrations did only slightly correlate with pH (R^2^ = 0.11) and were not associated with total bacteria grown (Figure S4A). The fraction of acetate showed strong negative correlations with both butyrate (R^2^ = 0.81) and propionate (R^2^ = 0.46) (Figure S4B, C), whereas relative concentrations of the latter two SCFAs were not associated (data not shown).

**Figure 3.**
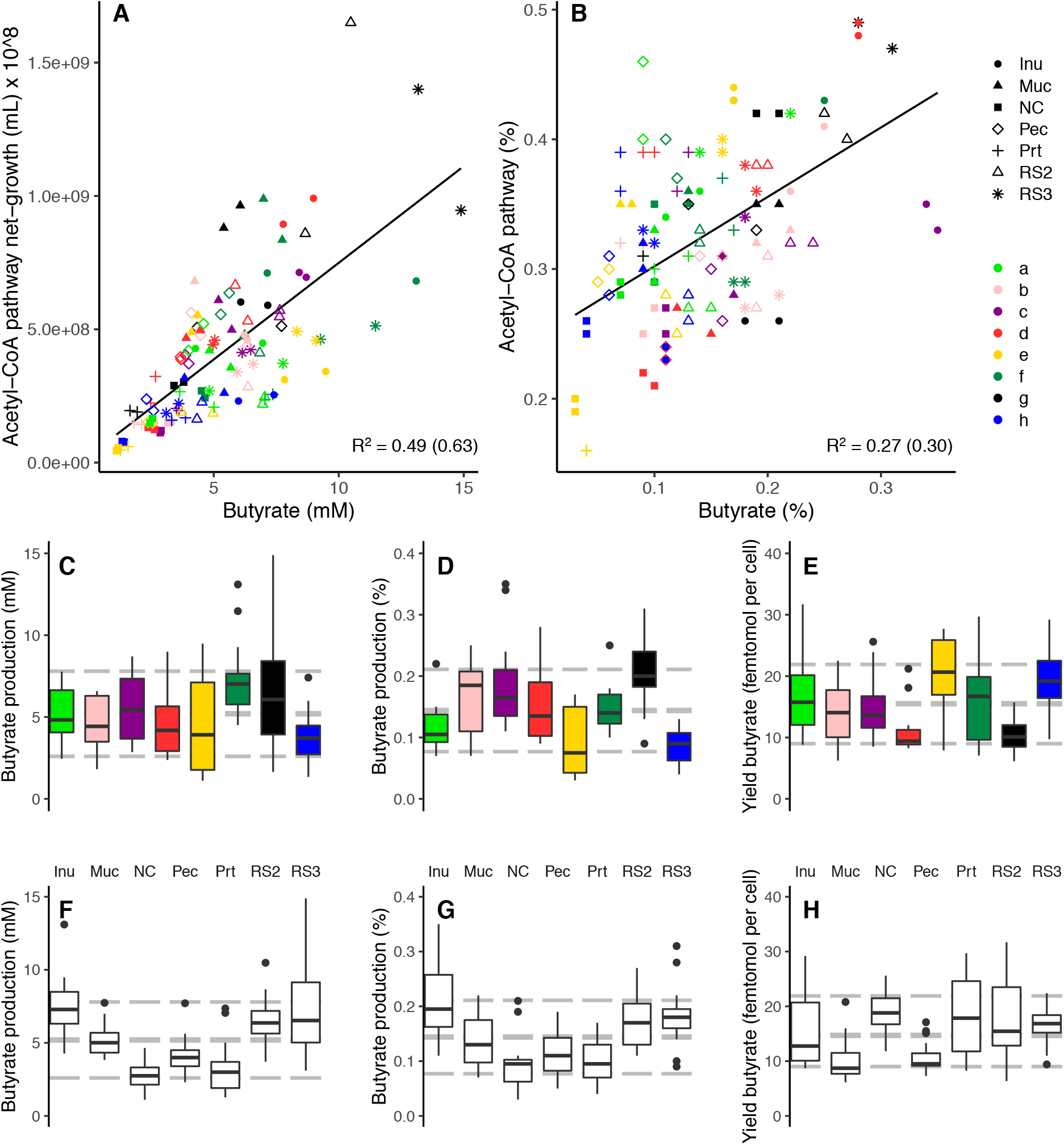
Correlation between ACoA pathway abundances and butyrate concentrations of *in vitro* experiments. Panel A shows the correlation between net-grown bacteria exhibiting the pathway and concentrations of butyrate formed, whereas associations between relative abundances of those bacteria with proportions of butyrate from total SCFAs are given in panel B. Values from communities derived from different donors and substrates are indicated. The Pearson correlation coefficient is given (values in brackets are based on log-transformed data). Panel C and F display concentrations of formed butyrate grouped into individual donors and substrates, respectively. Panel D and G gives corresponding results for relative butyrate concentrations (from total SCFAs), whereas panel E and H show respective yields, i.e., butyrate formed per grown bacterium harbouring the ACoA pathway. Grey lines depict average values along with standard deviations. Inu: inulin, Muc: mucin, NC: basal medium, Pec: pectin, Prt: protein, RS2/3: resistant starch type 2/3.

**Figure 4.**
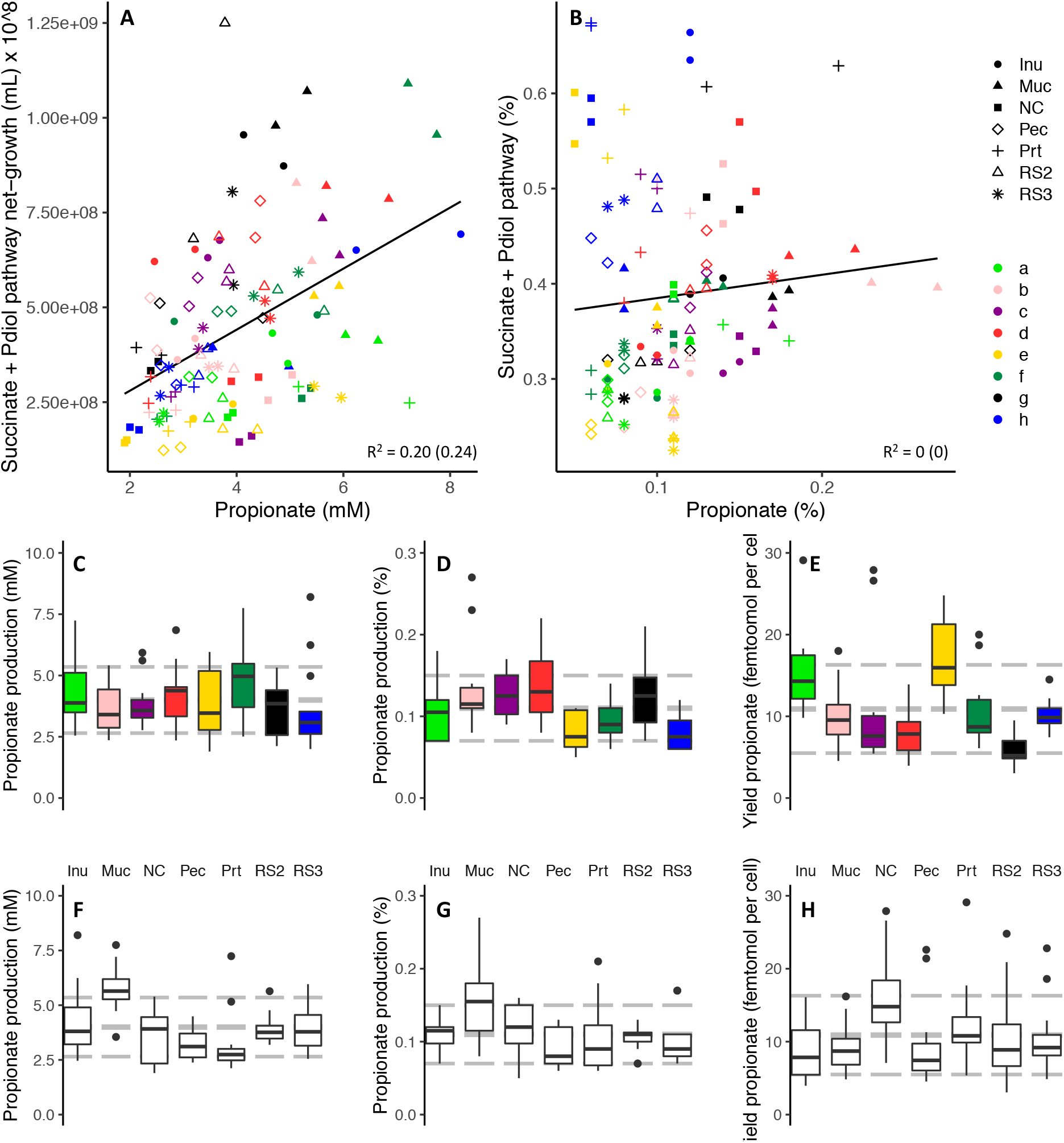
Correlation between propionate pathway abundances and propionate concentrations of *in vitro* experiments. Cumulative abundances of the Suc and Pdiol pathways are shown. Panel A shows the correlation between net-grown bacteria exhibiting those pathways and concentrations of propionate formed, whereas associations between relative abundances of those bacteria with proportions of propionate from total SCFAs are given in panel B. Values from communities derived from different donors and substrates are indicated. The Pearson correlation coefficient is given (values in brackets are based on log-transformed data). Panel C and F display concentrations of formed propionate grouped into individual donors and substrates, respectively. Panel D and G gives corresponding results for relative propionate concentrations (from total SCFAs), whereas panel E and H show respective yields, i.e., propionate formed per grown bacterium harbouring the Suc/Pdiol pathways. Grey lines depict average values along with standard deviations. Inu: inulin, Muc: mucin, NC: basal medium, Pec: pectin, Prt: protein, RS2/3: resistant starch type 2/3.

Predicted pathway abundances from 16S rRNA gene data correlated well with results derived from metagenomes displaying R^2^s of 0.60, 0.70 and 0.59 for the ACoA, Suc and Pdiol pathways, respectively (Figure S5A). Average abundances of pathways and composition of associated bacteria were also similar to metagenome-based data (Figure S5B, C).

### SCFA-producing communities *in vivo*

*In vitro* experiments above proved that it is possible to predict SCFAs production from sequence data, in particular in the case of butyrate. As a next step we investigated how those results relate to *in vivo* conditions by monitoring pathway abundances, bacterial concentrations and SCFAs concentrations in 20 individuals, who provided five fresh stool samples over a period of three months.

On average 28.1 % ± 5.5 % of bacteria carried the ACoA pathway, 14.4 % ± 7.5 % the Suc and 8.7 % ± 3.7 % the Pdiol pathways (Figure 5A); only 2.38 % of bacteria overlapped and carried the ACoA together with a propionate-forming pathway. Community composition of bacteria harbouring the ACoA pathway was in accordance with previously published data and that of *in vitro* results from above, where the bulk was classified as *Firmicutes* (95.9 %), with *Faecalibacterium* (19.3 %), *Agathobacter* (16.9 %) and Gemmiger (9.9 %) as the main taxa, and only a tiny fraction of *Bacteroidetes* (0.4 %) (Figure 5B). The Suc pathway exhibiting community was primarily composed of *Bacteroidetes* (68.7 %), with members of the genus *Bacteroides* (23.5 %) and *Phocaeicola* (20.1 %) representing the majority (Figure 5B), and of *Dialister* from the *Firmicutes* (20.0 %) (Figure 5B). Pdiol pathway carriers were almost exclusively of the *Firmicutes* (94.8 %), mainly of the genera *Blautia_A* (56.5 %) and *Anaerobutyricum* (21.7 %); the latter additionally exhibit the ACoA pathway representing the only noteworthy overlap between butyrate- and propionate-producers. Total as well as individual pathway community compositions showed strong subject-specific patterns (Figure S6).

**Figure 5.**
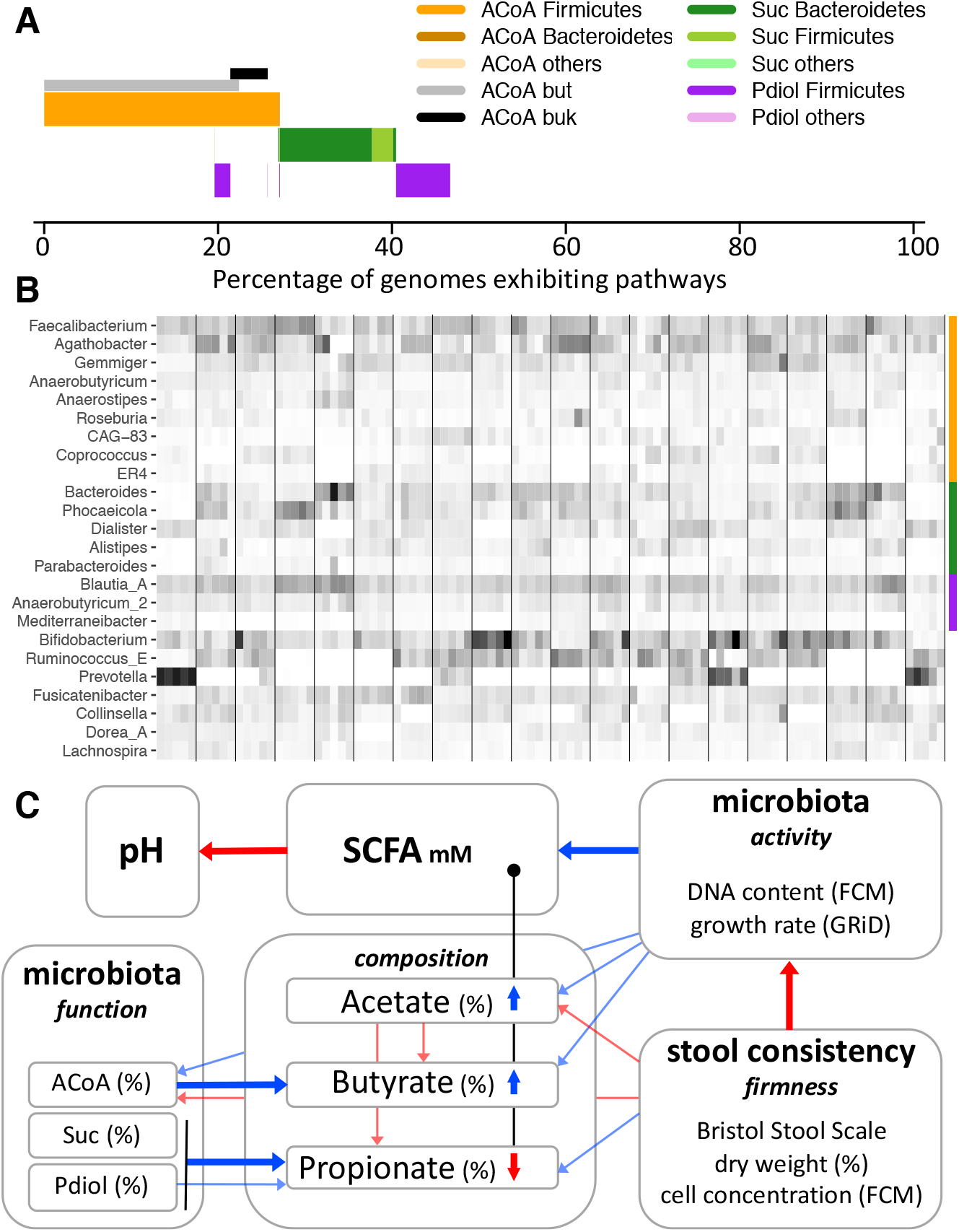
Overview of pathway abundances and taxonomic composition of *in vivo* experiment (20 subjects were sampled at five time-points over the period of three months). Panel A gives average abundances of pathways including taxonomic affiliations on the phylum level, whereas abundances of major genera comprising individual pathway communities (indicated by the colour bar on the right) are given in panel B; abundances are relative to house-keeping genes of the total bacterial community. In panel C, a mechanistic model on *in vivo* faecal SCFA concentrations based on individual parameters measured is given. Blue refers to positive correlations based on linear mixed-effect models that included subject as a random effect, whereas red depicts negative associations. Correlations that are considered most important are highlighted as thick arrows. For explanations see text.

Additional measured parameters from stool displayed strong variations between samples (Figure S7). SCFAs concentrations varied by an order of magnitude for acetate (17.9 - 164.1 mM; average 62.4 mM) and propionate (4.3 - 49.8 mM; average 21.0 mM), while butyrate concentration varied by a factor of forty (1.6 – 70.1 mM; average 18.8 mM). Bacterial concentrations ranged from 4.94×10^10^ to 5.98×10^11^ (average: 2.50×10^11^) cells per gram wet stool and from 2.77×10^11^ to 1.39×10^12^ (average: 8.84 ×10^11^) cells per gram dry faecal matter, respectively; faecal moisture content displayed wide variations (51.8 % – 85.4 %; average: 72.2 %). Values of the Bristol Stool Scale stretched over six of the seven categories (BSS 1 – 6).

Correlation analyses between all parameters allowed us to formulate a mechanistic model on factors governing faecal SCFAs concentrations that is shown in Figure 5C. All stool parameters displayed high subject-specific patterns (Figure S7) and we, hence, included subject as a random effect in our correlation analyses (individual results from generalized linear models are given in Table S1). Contrary to our expectations, no association between faecal SCFAs concentrations and the total amount of bacteria per gram stool was detected. However, bacterial activity, in particular green fluorescence signal intensities based on FCM analyses, which are a proxy of nucleic acid content, correlated positively with levels of faecal SCFAs (Figure 5C). However, this parameter was negatively correlated with stool firmness, which was measured as percent dry weight, stool texture according to the BSS and faecal cell concentrations that all correlated highly with each other (Table S1). The other parameter used to describe activity, namely, the growth rate index (GRiD) based on coverage ratios between *ori* and *ter* of constructed MAGs, correlated with FCM results and was trending (p ∼0.1) with total SCFAs concentrations. Stool firmness parameters did not correlate with total SCFAs concentrations. (Relative) butyrate concentrations were associated with both activity parameters as well as with total SCFAs concentrations, but not with stool firmness. However, firmness parameters correlated negatively with relative acetate concentrations and displayed positive associations with relative propionate concentrations (Figure 5C). The proportion of both butyrate and acetate were increased at higher total SCFA concentrations, while that of propionate was reduced. Relative acetate concentrations were negatively correlated with those of butyrate and propionate (Figure 5C). Abundances of ACoA, Pdiol and Suc pathways correlated with relative concentrations of respective SCFAs (Figure 5C). Firmness was negatively correlated with ACoA pathway carrying bacteria. A strong correlation between total SCFA concentrations and faecal pH was recorded, however, pH was not associated with abundances of any pathways (Table S1).

Predicted pathway abundances from 16S rRNA gene data were similar as those based on metagenomic analyses displaying R^2^s of 0.67, 0.80 and 0.52 for the ACoA, Suc and Pdiol pathways, respectively (Figure S8A); also the overall composition was comparable between the two techniques (Figure S8B). Detected average abundances of Suc were similar between the two methods, whereas concentrations of the ACoA and Pdiol pathways were higher compared with metagenomic results (Figure S8C).

### Temporal stability of pathway abundances and SCFA concentrations

The longitudinal character of our study enabled insights into temporal dynamics of SCFAs concentrations and bacteria harbouring pathways for their formation. We observed a high volatility of butyrate concentrations displaying 41.2 % ± 22.8 % average difference between the first time-point one and all other time-points (Figure 6A). Fluctuations in its relative concentration was much less (20.7 % ± 15.7 %) (Figure 6B), which was in accordance with relative abundances of ACoA pathway carriers that were rather constant varying on average by only 15.1 % ± 11.6 % (Figure 6C). Propionate pathway carriers showed higher dynamics for both the Suc (22.3 % ± 16.6 %) and the Pdiol (23.5 % ± 17.7 %) pathway (Figure 6C). Measured concentrations of propionate were, however, less volatile (31.2 % ± 19.3 %) as those of butyrate, as were relative propionate concentrations (17.7 % ± 13.4 %) (Figure 6A, B). Acetate showed highest temporal stability for both absolute (29.5 % ± 19.5 %) and relative concentrations (8.0 % ± 7.4 %).

**Figure 6.**
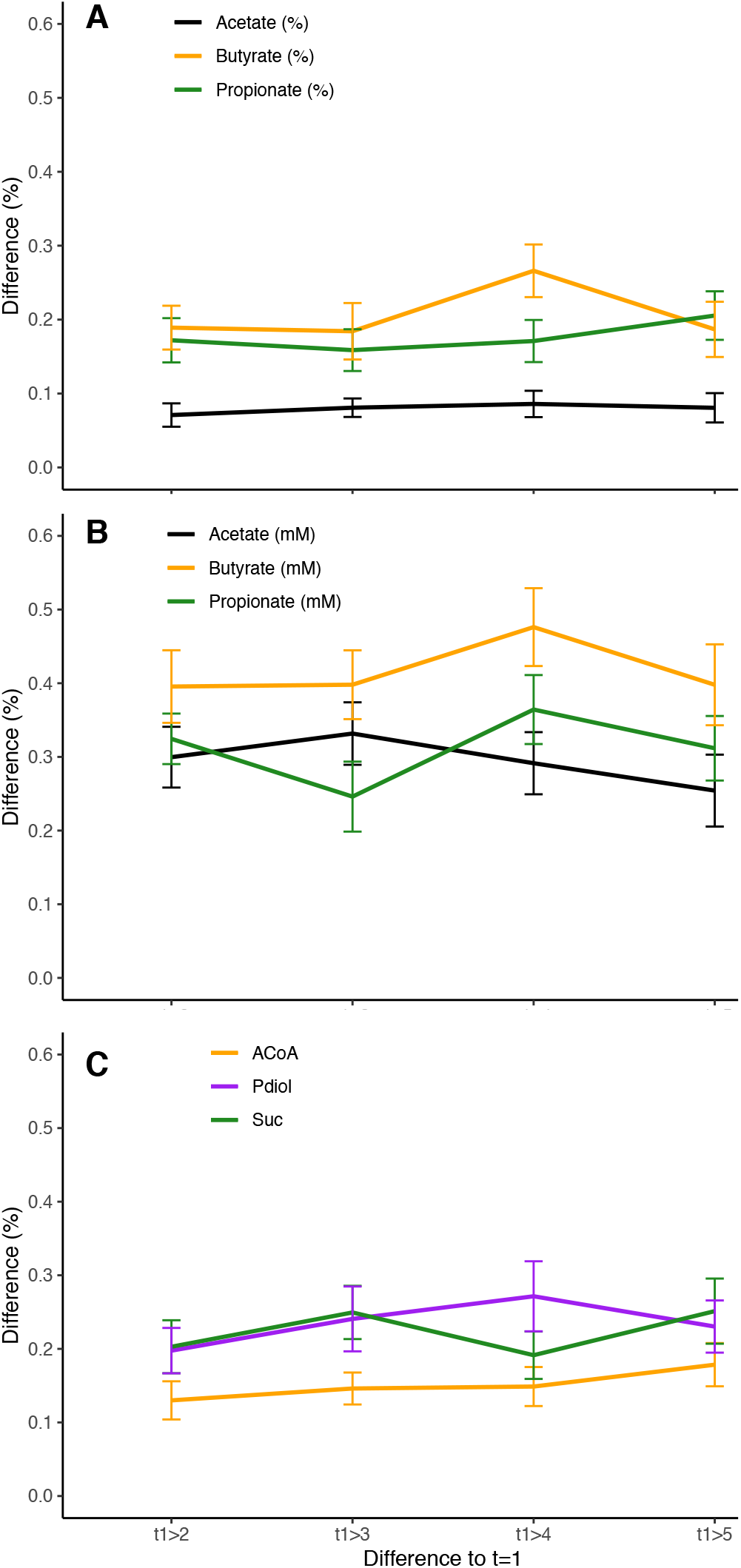
Temporal stability of SCFA concentrations and pathway abundances *in vivo*. Subjects (n=20) were sampled (n=5) over a period of three months and results relative to the first time-point are shown. Panel A displays relative concentration changes of the SCFAs acetate, butyrate and propionate, whereas variations based on proportions (relative to total SCFA concentrations) and of abundances of bacteria exhibiting individual pathways are given in panel B and C, respectively.

## Discussion

The aim of this study was to gain quantitative insights into functional communities that form butyrate and propionate, and to predict the production of those SCFAs based on sequence data derived from both metagenomes and the 16S rRNA gene. Several criteria have to be fulfilled in order to achieve those goals. 1.) Accurate databases of gut bacteria carrying SCFAs synthesis pathways are required, 2.) pathways should not co-occur on genomes excluding metabolic flexibility for production of the two SCFAs, 3.) for predictions based on 16S rRNA gene data pathways have to be distributed following phylogenetic patterns, 4.) yields, i.e., amount of SCFAs produced per cell harboring respective pathways, should be equal between taxa and 5.) should not be governed by environmental factors (e.g. type of growth substrate).

We comprehensively screened for pathway abundances in isolates and MAGs specifically derived from the gut environment provided by the UHGG (23). A manually curated database for butyrate-producing bacteria was already established previously (16), but no systematic genome screening for propionate-forming pathways has been performed so far. On a genome level, our results do largely agree with KEGG, representing one of the most widely used public database, however, several crucial discrepancies were revealed. Most importantly, many *Enterobacteriaceae* were wrongly predicted to have the ACoA pathway by KEGG and *Akkermansia*, as well as several *Veillonella*, that are known propionate-producers (34)(15), were suggested to lack this function by KEGG. Furthermore, no information on terminal enzymes of the ACoA pathway, namely butyryl-CoA:acetate CoA transferase (*but*) and butyrate kinase (*buk*), are available within KEGG. Specifically for the ACoA pathway it is important to consider pathway completeness, because a multitude of mis-annotations based on gene-homology alone are present (14)(35). Those points highlight the use of manually-curated databases for specific functions of interest.

The largely coherent distribution of the ACoA and Suc pathway on the genus level suggests strong selection for those features and strengthens the view that both pathways serve as core fermentative routes in respective bacteria (15, 16). We have previously demonstrated that taxonomy-based approaches (on the genus level) for predicting ACoA pathway abundances is valuable (16). Nevertheless, our analyses here suggested that within a few genera, such as *Blautia* and *Eubacterium*, this metabolic route is not homogenously present. However, it still follows phylogenetic clustering, making phylogenetic-based predictions superior over plain taxonomic-based analyses, as suggested earlier (36)(37). This is also true in the case of propionate, where the Pdiol pathway displayed phylogenetic patterns within certain genera, where not all members exhibited the pathway (e.g. *Mediterraneibacter, Eisenbergiella*). Our data further revealed a profound separation between bacteria forming either butyrate or propionate. *In vivo* results demonstrated that a tiny part (2.38 %) of bacteria cannot be unequivocally assigned either group. Bacteria harbouring the genetic make-up for both functions switch their metabolism according to the prevailing environmental conditions hindering predictions on SCFAs profiles on the DNA level. For instance, *R. inulinivorans* forms propionate during growth on fucose due to substrate-induced gene expression instead of butyrate, which is the primary SCFAs formed when growing on glucose (38). Protein-fed butyrate production routes have not been considered here, as they are not vital for growth of individual bacterial carriers and abundances are, hence, uncoupled from activity (16). Furthermore, proteins are believed to play only minor roles as growth substrates in the large bowel and their conversion to butyrate can also be catalysed by the ACoA pathway making this pathway the primary route for butyrate formation from proteins in the colonic environment (13). In summary, the first three criteria mentioned above are fulfilled and form the basis for predicting butyrate- and propionate-forming communities along with metabolite concentrations from sequencing data.

An additional crucial aspect for predicting SCFAs production is functional redundancy, where pathway activity has to be constant between taxa and environmental conditions (e.g. growth substrates). Our *in vitro* experiments illustrated that yields were in a similar range, regardless of the type of substrate supplied and of individual’s community composition. In other words, the amounts of butyrate and propionate produced per grown bacterium harbouring a pathway were very similar, demonstrating functional redundancy of bacteria and independence of growth substrates. Those results are in-line with previous observations (19). However, exact values on yields of whole functional communities have not been reported so far, as their determination requires adequate methodologies for enumerating pathway-carrying bacteria that was achieved here by coupling enumeration of bacteria via FCM with metagenomic. Furthermore, the experimental set-up has to be designed in order to assure growth, i.e., multiplication, of bacteria. In our experiments we specifically diluted starting communities to provide growth over two orders of magnitudes (from ∼10^7^ mL^−1^ to ∼10^9^ mL^−1^). Often, communities are incubated at high cell concentrations with relatively little amount of substrates, which works well for assessing production capabilities for individual SCFAs (39), but hampers accurate calculations of yields. It should be noted that yields do not refer to absolute final cell growth. We did observe substantial differences between substrates in terms of SCFA concentrations/compositions and abundances of functional communities. For instance, the resistant starches were confirmed to promote formation of butyrate (and bacteria containing the ACoA pathway) (40)(41), which was also the case for inulin (42). On the other hand, mucin was rather promoting formation of propionate, which is in accordance with major mucin-degrading taxa, such as specific *Bacteroides*, exhibiting the Suc pathway (43).

Complying with all five requirements introduced above, our faecal incubation experiments have demonstrated that *in vitro* it is indeed possible to calculate the absolute production of butyrate, and its proportion of the total SCFA pool, based on enumerating bacteria that exhibit the ACoA pathway. For propionate, predictions on absolute concentrations were possible, however, with less accuracy, and the fraction of propionate from total SCFAs were not explainable based on pathway abundances *in vitro* (*in vivo* they did, however, correlate). We do attribute this observation rather to physiology of those bacteria than to inaccurate pathway callings. For instance, the Pdiol pathway is considered to be a major route of metabolic cross-feeding taking lactate produced by other bacteria as input (15), which is not essential for growth of those bacteria. *B. obeum*, for instance, can grow on sugars without producing any propionate (44), and specific induction of genes from this pathway was shown to depend on the growth substrate, as discussed above (38). Nevertheless, calculating propionate concentrations from the Suc pathway abundance alone was less effective and we, hence, used cumulative pathway abundances in our analyses. For bacteria exhibiting the Suc pathway physiological adaptations were reported as well, where rather succinate and acetate than propionate are produced under certain conditions (44). The former compound is an intermediate and does usually not accumulate in the SCFAs pool of gut communities (13)(19).

*In vivo*, relative abundances of the ACoA pathway and of the two propionate-forming routes did correlate with relative concentrations (proportions) of the two SCFAs, demonstrating that functional communities are reflected in SCFAs composition. Against our expectations, total SCFAs concentration did not correlate with absolute abundance of bacteria, nor did we observe an association between absolute abundances of pathways and corresponding SCFAs concentration. Results suggest that bacterial concentration is merely governed by stool firmness that is directly connected with retention time (45). The longer the colonic transit time of faecal matter the more moisture is absorbed resulting in a higher dry weight and higher bacterial concentrations per gram of stool. Water, ions and SCFAs are absorbed alike decoupling bacterial concentrations from SCFAs concentrations (46). Furthermore, it can be assumed that conditions comprising less water content caused by slower transit provide challenging environments for bacterial growth, which is mirrored by the negative correlation observed between microbial activity and stool firmness, further decoupling SCFAs concentrations from cell numbers. Community structure (47) and functionality (48) was previously associated with stool consistency, which is supported here, where longer transit (firmness) was negatively associated with ACoA pathway abundance and an enrichment for propionate production. According to our calculations based on *in vitro* yields, the amounts of SCFAs absorbed by the host are 97.7 % % ± 2.2 % and 94.5 % % ± 5.4 % for butyrate and propionate, respectively, which is even higher than previously suspected (11, 12), and stresses the distinction between SCFAs production (defined as the total amount of SCFAs formed in a defined period of time) and SCFAs concentrations. The latter can be regarded as a snapshot parameter, which is highly influenced by a series of factors irrespective of the source, i.e., bacterial concentrations, which was also reflected in high temporal volatilities. On the other hand, proportions of individual SCFAs were rather constant, as were corresponding pathway abundances, and variances of neither parameters correlated with time intervals between sampling points, indicating that individual’s SCFAs pattern, and corresponding functional communities, respectively, are fairly stable over time.

In conclusion, we give detailed insights into butyrate- and propionate-producing communities revealing that they form two taxonomically distinct groups in gut microbiota, whose abundances determine SCFAs concentrations *in vivo*. Overall, it was possible to predict relative metabolite concentrations from bacteria carrying respective pathways, demonstrating that altering the community structure is a valuable strategy to promote production of those specific SCFAs. The successful use of the 16S rRNA gene for function prediction provides a high-throughput, low-cost screening alternative over more tedious metagenomic analyses, which facilitates investigations on SCFAs-forming communities in broad-scale applications.

## Acknowledgements

We greatly thank all our participants contributing to the *in vivo* experiment. Furthermore, thanks to Maren Scharfe and Michael Jarek from the HZI for their technical assistance and to Colin Davenport and Jannes Gless for maintaining the HPCSeq at MHH. This work was funded by the DFG (project #456214861) and intramural funds (HiLF II of MHH).

